# Seed morphological traits as a tool to quantify variation maintained in *ex situ* collections: a case study in *Pinus torreyana* (Parry)

**DOI:** 10.1101/2020.09.18.303768

**Authors:** Lionel N Di Santo, Monica Polgar, Storm Nies, Paul Hodgkiss, Courtney A Canning, Jessica W Wright, Jill A Hamilton

## Abstract

**Background:** Understanding the within- and among-population distribution of trait variation within seed collections may provide a means to approximate standing genetic variation and inform plant conservation.

**Aims:** This study aimed to estimate population- and family-level seed trait variability for existing seed collections of Torrey pine (*Pinus torreyana*), and to use these data to guide sampling of future collections.

**Methods:** We quantified variation in 14 seed morphological traits and seedling emergence within and among Torrey pine populations. Using a simulation-based approach, we used estimates of within-population variance to assess the number of maternal families required to capture 95% of trait variation within each existing seed collection.

**Results:** Substantial structure was observed both within and among Torrey pine populations, with island and mainland seeds varying in seed size and seed coat thickness. Despite morphological differences, seedling emergence was similar across populations. Simulations revealed that 83% and 71% of all maternal families within island and mainland seed collections respectively needed to be resampled to capture 95% of seed trait variation within existing collections.

**Conclusions:** From a conservation perspective, our results indicate that to optimize genetic diversity captured in Torrey pine seed collections, maximizing the number of maternal families sampled within each population will be necessary.

## Introduction

Ex situ seed collections preserve species genetic diversity outside of their native range, providing the raw material for species reintroductions and germplasm to augment restoration (Guerrant Jr *et al*. 2014; Potter *et al*. 2017). Ensuring ex situ collections represent genetic variation found in natural populations is critical to both contemporary conservation and potential future restoration efforts (Schaal and Leverich 2004; Basey *et al*. 2015). An invaluable conservation resource, particularly for rare species, ex situ collections protect against biodiversity loss in the wild, while preserving species’ evolutionary potential. However, the cost and logistical constraints associated with seed collection pose a significant challenge. Given this challenge, means are needed to optimize ex situ sampling efforts (Hoban and Schlarbaum 2014; Di Santo and Hamilton 2020).

One approach may be to use the distribution of trait variation existing within contemporary *ex situ* seed collections as a proxy for quantifying standing genetic variation within and among populations. Although multiple factors influence plant phenotypes (Monty *et al*. 2013; Villellas *et al*. 2014), seed morphological variation is often considered highly heritable. For example, seed length, seed width, and seed mass have a heritability (or repeatability) estimated between 0.33 and 0.98 in conifers, including maritime pine (*Pinus pinaster*), chir pine (*Pinus roxburghii*), and white spruce (*Picea glauca*) (Roy *et al*. 2004; Carles *et al*. 2009; Zas and Sampedro 2015). In addition, traits such as seed shape, seed coat thickness, or embryo length also exhibit high heritability, with values estimated between 0.59 and 0.96 for agronomic species, including soybean (*Glycine max*), narrow-leafed lupin (*Lupinus angustifolius*) and rice (*Oryza sativa*) (Pandey *et al*. 1994; Cober *et al*. 1997; Mera *et al*. 2004; Hakim and Suyamto 2017). Given these observations, variation in seed morphological traits likely has a genetic basis and may reflect standing genetic variation within and among populations. In addition, morphological variation of seeds stored ex situ may reflect variation attributable to the maternal environment (Platenkamp and Shaw 1993; Singh *et al*. 2017). However, for rare species where existing genetic data are limited, quantifying within and between population variation for traits largely considered heritable within existing seed collections may be invaluable to optimizing future collections, even if estimates of genetic variation do not control for maternal environment.

The distribution of heritable genetic variation estimated via common garden experiments – experimental approaches used to understand the genetic contribution to phenotypic variation under common environmental conditions – (Weber and Kolb 2014; Hamilton *et al*. 2017; Yoko *et al*. 2020) or molecular genetic data (Zhang and Zhou 2013; Hausman *et al*. 2014; Tamaki *et al*. 2018) can be used to quantify the distribution of standing genetic variation. However, when common garden experiments or molecular genetic data are unavailable, quantifying trait variation within and among ex situ seed population collections may provide a reasonable proxy for the distribution of genetic variation. Millions of seed accessions have been stored in gene banks internationally (FAO 2010), representing a large conservation and research resource. Although common garden experiments are preferred when available, heritability of seed morphological traits and ease of access to seeds through existing ex situ collections suggests that quantifying seed morphological variation may provide a timely approach to estimating variation preserved in collections. In addition, where the goal is to limit the loss of biodiversity and preserve evolutionary potential for rare species, existing seed morphological data may be leveraged to optimize supplemental conservation collections.

*Pinus torreyana* Parry (Torrey pine), is one of the rarest pines in the world (Critchfield and Little 1966; Dusek 1985), endemic to two discrete natural populations in California. Torrey pine occupies one mainland population (*Pinus torreyana* subsp. *torreyana*) of approximately 6,000 trees at the Torrey Pine State Reserve in La Jolla, CA, and an island population (*Pinus torreyana* subsp. *insularis*) of approximately 3,000 reproductively mature trees on Santa Rosa Island, CA, one of the Channel Islands (Ledig and Conkle 1983; Haller 1986; Hamilton *et al*. 2017) (Fig. 1). Listed as critically endangered by the IUCN (2020), Torrey pine is of critical conservation concern due to multiple factors, including low population size (Franklin and Santos 2011; Hall and Brinkman 2015), low genetic diversity (Ledig and Conkle 1983; Whittall *et al*. 2010), climate change, and environmental and human-mediated disturbances (Franklin and Santos 2011; Hamilton *et al*. 2017). While in situ conservation has preserved the whole of the species’ range, with fewer than 10,000 reproductively mature individuals in native populations, there are substantial risks for population-level extirpation events. To mitigate potential losses in the wild, conservation efforts have focused on preservation of seed ex situ. While ex situ seed collections provide an invaluable conservation resource, they may also be used to quantify species’ trait variation needed to inform future conservation efforts.

**Fig. 1.**
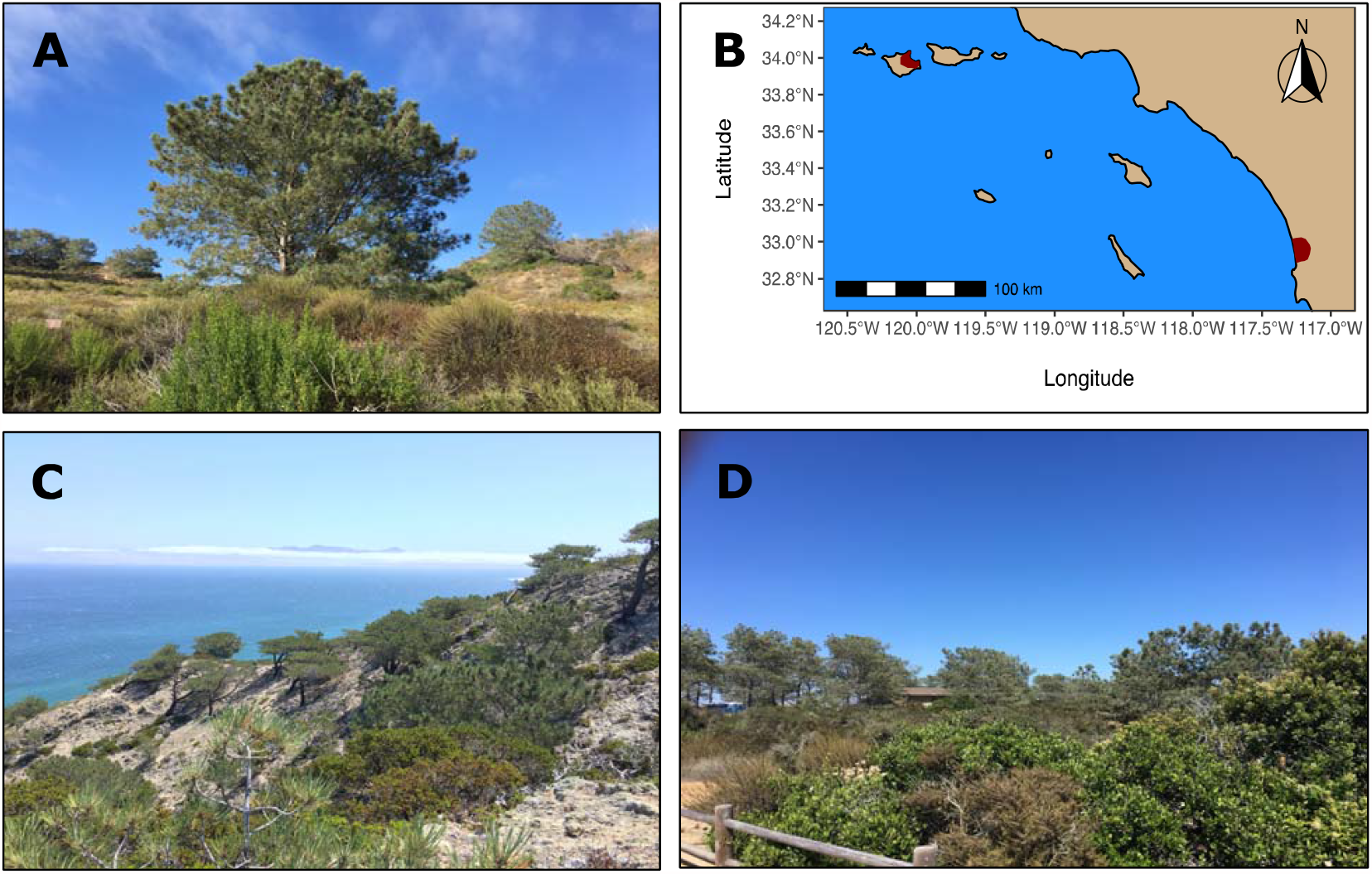
(A) *Pinus torreyana* individual. (B) *Pinus torreyana* distribution map, including Torrey pine distribution on Santa Rosa Island, CA (*Pinus torreyana* subsp. *Insularis*, top left red-shaded area) and at the Torrey Pine State Reserve, CA (*Pinus torreyana* subsp. *torreyana*, bottom right red-shaded area). (C) Torrey pine stand on Santa Rosa Island, CA. (D) Torrey pine stand at the Torrey Pine State Reserve, CA.

In this study, we evaluate morphological trait variation in a large ex situ conservation collection of Torrey pine seed sourced from the two native extant populations. Specifically, we quantify the distribution of variation for 14 seed morphology traits and assess differences in emergence between island and mainland seedlings. In addition, we use existing ex situ collection data to provide supplemental population sampling guidance for future Torrey pine collections. For this latter objective, we use simulations to estimate the number of maternal families required to capture 95% of seed morphological variation existing in contemporary ex situ collections, for both island and mainland population independently. This study evaluates the distribution of seed morphological variation in ex situ collections as a proxy for standing genetic diversity, quantifying variation attributable to within and between population differences. These data are then used to inform population sampling necessary to meet conservation objectives in future seed collections. Although presented here using Torrey pine, our approach is broadly applicable for ex situ collections within species with largely heritable seed trait variation.

## Materials and methods

### Cone collection and seed processing

Mature, open-pollinated Torrey pine (*Pinus torreyana* Parry) cones were collected from native extant populations as part of a large ex situ conservation collection between June and July of 2017. Cones were collected from 157 trees on Santa Rosa Island (Channel Islands National Park), CA (island population) and 201 trees at the Torrey Pine State Reserve in La Jolla, CA (mainland population), representing the species’ entire natural distribution (Fig. 1; See Supporting Information Figure S1). Where possible, we collected between five to ten cones per maternal tree at each location. Sampling of reproductive maternal trees was evenly spaced; however, adjacent maternal trees were occasionally sampled to ensure enough cones were collected. On average, mainland and island trees sampled were separated by approximately 714 (range = 0 – 2,092) and 397 (range = 1 – 1,131) meters, respectively. Seeds, organized by individual maternal tree, were then extracted from cones using a combination of mallet and pliers and processed for inclusion in a long-term ex situ conservation collection (see below).

### Seed viability tests

Estimating viability of seeds preserved *ex situ* is necessary given their potential role in restoration, reforestation, or reintroduction. Given this, the potential viability of Torrey pine seeds was tested using two complementary approaches prior to inclusion in the final ex situ collection. A float test was first used as a rapid, low-cost approach to assess seed viability. Floating seeds were presumed to lack an endosperm or embryo, while seeds that sunk were presumed filled. Seeds were dropped into water for approximately 15 seconds to differentiate presumed non-viable, floating seeds from presumed viable, sinking seeds (Gribko and Jones 1995; Morina *et al*. 2017). Those seeds classed as likely viable were organized by maternal tree using paper bags, and then placed in a Blue M drying oven (Thermal Product Solutions, White Deer, Pennsylvania, USA) maintained at 37°C for 24 hours to remove potential surface moisture. Following this, seeds from a haphazard sample of maternal families were x-rayed at the Placerville Nursery, CA. In addition to visualizing seed morphological variation, x-ray photographs were used to verify viability based on float tests. Acrylic seed trays [20.3 cm x 25.4 cm x 0.48 cm], with a 9 × 11 array of wells, were used to separate and position each Torrey pine seed over the x-ray film. Kodak x-OMAT HBT film (20.3 cm × 25.4 cm) was placed in a lightproof x-ray film cassette which was positioned in the x-ray machine with the seed tray centered on top of the film, with a shelf height of 55.9 cm. The x-ray was taken using a 17 kVP exposure for a total of two minutes, based on standardized conditions established previously for *Pinus coulteri* (Sara Wilson, USDA Forest Service, pers. comm.). X-ray images were digitized using a Nikon D40 digital camera mounted on a tripod over a light box.

### Morphological measurement of seed traits

Using ImageJ (Abràmoff *et al*. 2004), eight seed morphological traits were measured across 80 mainland maternal families and 30 island maternal families, representing a haphazard subset of the complete collection (Fig. 2; Table 1). Each x-ray picture was scaled using the diameter of a seed tray well (1.87 cm) to express pixels as trait values in centimeters. Directly measured seed traits included seed length (SL, cm), seed width (SW, cm), embryo length (EL, cm), embryo width (EW, cm), seed coat width (SCW, cm), seed area (SA, cm^2^), endosperm area (ESA, cm^2^), and embryo area (EA, cm^2^). We selected these traits as they can readily be measured from x-ray pictures of seeds and provide a ubiquitous means to evaluate morphological variation for plants preserved ex situ. Using measured seed traits, six additional traits were derived (Table 1), including seed length/width ratio (SLW), embryo length/width ratio (ELW), relative embryo size (RES), relative endosperm size (REndS), seed coat area (SCA, cm^2^), and relative seed coat size (RSCS). These traits were derived as they provide a means to relate different morphological traits to each other and can provide a fine-scale estimate of the relative contribution of growth and size traits within individual seeds. We measured five randomly selected seeds per maternal tree, including three technical replicates per seed for each trait (the same seed was measured three times for any given morphological trait). Measurements were averaged across technical replicates to summarize the mean trait value per seed. In total, 550 seeds were measured from across 110 maternal trees spanning the two Torrey pine populations.

**Table 1.** Morphological measurements of Torrey pine seeds sourced from Santa Rosa Island (Island, n=30) and Torrey Pine State Reserve (Mainland, n=80), CA. Listed are population mean estimates (±SE) of measured (A) and derived (B) seed traits summarized by maternal families. Measurable traits: seed length (SL), seed width (SW), embryo length (EL), embryo width (EW), seed coat width (SCW), seed area (SA), endosperm area (ESA), and embryo area (EA). Derived traits: seed length/width ratio (SLW), embryo length/width ratio (ELW), relative embryo size (RES), relative endosperm size (REndS), seed coat area (SCA), and relative seed coat size (RSCS). Differences in seed morphology between mainland and island populations were significant (α=0.05) for all 14 seed traits.

| Population | SL (cm) | SW (cm) | SCW (cm) | EL (cm) | EW (cm) | SA (cm<sup>2</sup>) | ESA (cm<sup>2</sup>) | EA (cm<sup>2</sup>) |
| --- | --- | --- | --- | --- | --- | --- | --- | --- |
| Mainland | 1.41±0.016 | 0.77±0.009 | 0.09±0.002 | 1.08±0.012 | 0.15±0.003 | 0.89±0.023 | 0.55±0.014 | 0.17±0.004 |
| Island | 1.67±0.02 | 0.97±0.015 | 0.12±0.002 | 1.2±0.02 | 0.19±0.004 | 1.41±0.031 | 0.81±0.021 | 0.23±0.007 |

**B – Derived seed traits**
| Population | SLW<br>(SL/SW) | ELW<br>(EL/EW) | SCA (cm<sup>2</sup>)<br>(SA-ESA) | RES<br>(EA/ESA) | REndS<br>(ESA/SA) | RSCS<br>(SCA/SA) |
| --- | --- | --- | --- | --- | --- | --- |
| Mainland | 1.86±0.013 | 7.28±0.094 | 0.34±0.01 | 0.31±0.005 | 0.62±0.003 | 0.38±0.003 |
| Island | 1.77±0.037 | 6.64±0.136 | 0.6±0.013 | 0.29±0.004 | 0.57±0.004 | 0.43±0.004 |

**Fig. 2.**
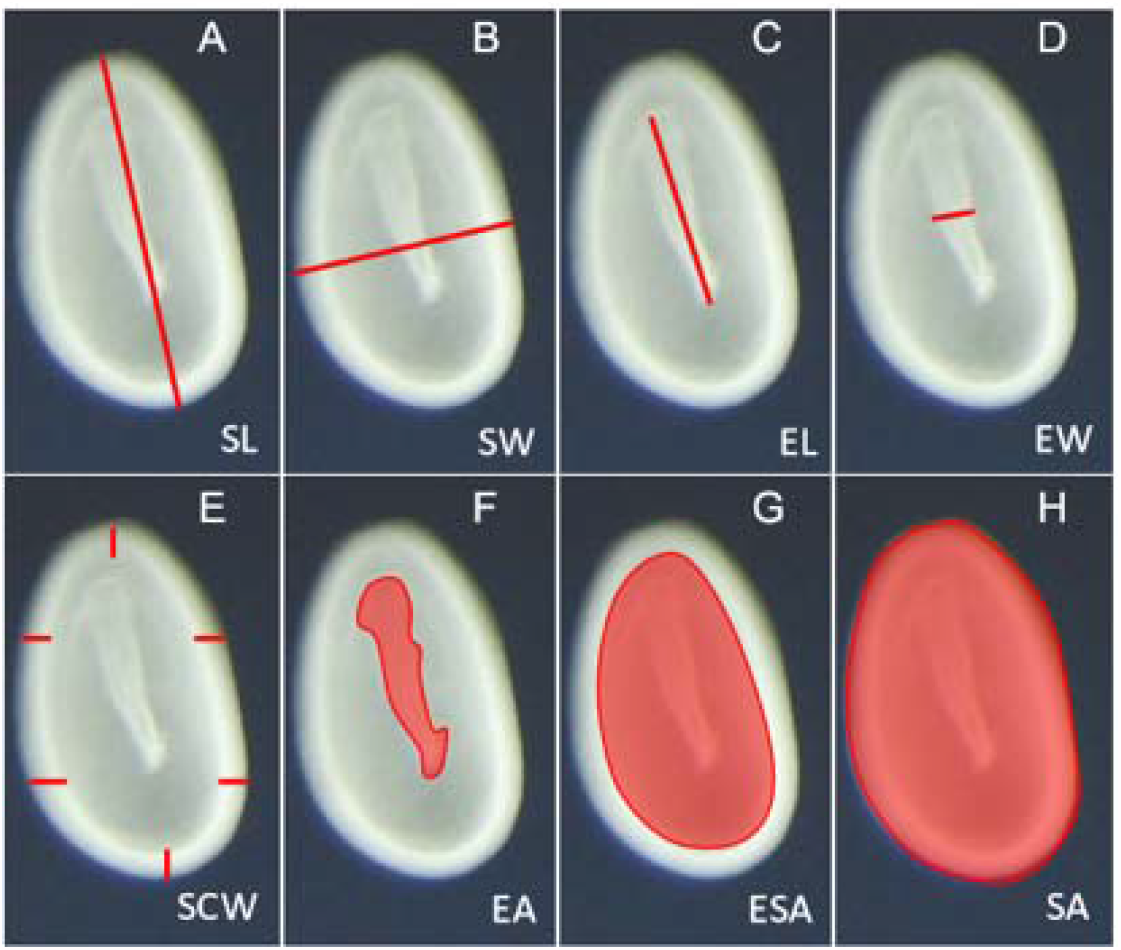
Visual of morphological measurements taken using ImageJ for seeds collected on Santa Rosa Island and at the Torrey pine State Reserve. (A) Seed length [cm]. (B) Seed width [cm]. (C) Embryo length [cm]. (D) Embryo width [cm]. (E) Seed coat width [cm]. (F) Embryo area [cm^2^]. (G) Endosperm area [cm^2^]. (H) Seed area [cm^2^].

### Seedling emergence test

Within a restoration or reintroduction context, concurrent seedling emergence is often preferred for nursery plantings. To evaluate the timing and probability of seedling emergence within Torrey pine populations, a trial was conducted in January 2018 using a random subset of seeds from the ex situ collection, including seeds sourced from Torrey Pine State Reserve and Santa Rosa Island, CA. Following x-ray, seeds were stored at 4°C in sealed mylar bags (USA emergency supply, Beaverton, Oregon, USA) placed in plastic boxes; each box contained desiccant crystals to decrease ambient moisture and reduce likelihood of mold. Seeds from eight maternal families per population were selected for the emergence trial. Between eight to ten seeds per maternal tree were weighed and then stratified under cold, moist conditions for 30 days (placed in plastic boxes on a moist paper towel at 4°C). Seeds were sown directly into a 164 mL Ray Leach “Cone-tainer” ™ (Stuewe & Sons, Tangent, Oregon, USA) filled with Sunshine® Mix #4 (Sungro horticulture, Agawam, Massachusetts, USA), pressed halfway into the soil, and then covered with a thin layer of gravel. For approximately one month following planting, seeds were misted for one minute at hourly intervals over a daily eight-hour period (9am – 4pm). Following emergence, seedlings were hand watered to saturation weekly to biweekly as needed. Emergence was quantified across three separate timepoints (Feb 06/2018, Feb 16/2018, and Mar 07/2018) per maternal family as the proportion of seeds that successfully developed into living seedlings from the total initially planted.

### Evaluating the distribution of seed trait variation

We conducted a principal component analysis (PCA) using all 14 measured and derived seed traits averaged by maternal family to evaluate population-specific differentiation in seed morphology. Prior to performing the PCA, to account for differences in measurement units, all seed traits were standardized using the *scale()* function in R implementing the z-score standardization: 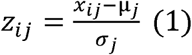, where x_*ij*_ is the non-transformed trait value, μ_*j*_ is the mean of a given seed trait across populations, and a the standard deviation of the same seed trait across populations. Subsequently, to test for seed trait differences between population means, we used either Student’s two-sample test or its non-parametric equivalent when normality was not met, Wilcoxon’s two-sample test, within the R package “exactRankTests” (Hothorn and Hornik 2019). Normality was assessed using Shapiro-Wilk’s test of normality within each population. In total, four of the fourteen traits were distributed normally in both Torrey pine populations, including seed width (mainland: W = 0.97, P = 0.06; island: W = 0.97, P = 0.52), embryo length (mainland: W = 0.98, P = 0.29; island: W = 0.95, P = 0.21), embryo width (mainland: W = 0.97, P = 0.09; island: W = 0.93, P = 0.05), and embryo area (mainland: W = 0.98, P = 0.45; island: W = 0.96, P = 0.26).

To evaluate the distribution of morphological trait variation within and between Torrey pine populations, we quantified the proportion of variation attributed to population and maternal tree families using measured and derived morphological traits summarized by seeds. For each trait, we fit a linear mixed model using the R package “lme4” (Bates *et al*. 2015) with population considered a fixed effect and maternal families within populations considered a nested random effect: *Y*_*ij*_ =μ + *π*_*j*_ + *r*_*i*/*j*_ + *e*_*ij*_, where *Y* is the observed seed trait value, μ is the seed trait overall mean, *π*_*j*_ is the effect of population origin on the seed trait mean, *r*_*i*/ *j*_ is the effect of maternal family within populations on the observed seed trait value, and e_*ij*_ are the effects on the seed trait value of any other variables unaccounted for in the model (residual error). For each model, normality of residual errors was visually assessed and significance of fixed- and random-effect terms was tested using the functions *anova()* and *ranova()* respectively, implemented in the R package “stats” (R Core Team 2020) and “lmerTest” (Kuznetsova *et al*. 2017). Proportions of seed morphological variance explained by populations (marginal R^2^, R^2^_m_), both populations and maternal families (conditional R^2^, R^2^ _c_), and maternal families alone (R^2^_c_ - R^2^ _m_) were quantified for each model independently using the function *r*.*squaredGLMM()* implemented in the R package “MuMIn” (Bartoń 2020).

### Assessing differences in seedling emergence across populations

To test for differences in the probability and the timing of seedling emergence in Torrey pine, we evaluated the proportion of seeds that produced seedlings both within and between populations across timepoints. First, we used Friedman’s rank sum test (non-parametric repeated measures ANOVA) followed by Wilcoxon’ paired two-sample test, both implemented in the R package “rstatix” (Kassambara 2020), to assess differences in the proportion of emerged seedlings between timepoints within populations. We used a non-parametric approach for both Torrey pine populations because normality could not be assumed at select timepoints due to high frequency of zero values. We accounted for multiple testing using Benjamini and Hochberg (1995)’s False Discovery Rate (FDR) correction implemented in the *wilcox_test()* function. Following this, we evaluated timepoint-specific population differences in seedling emergence. We used Shapiro-Wilk’s test to assess populations’ deviation from normality at each timepoint and either Student’s (for timepoints passing the normality test) or Wilcoxon’s two-sample test (for timepoints failing the normality test) to evaluate differences in population emergence. Timepoints Feb 16/2018 (mainland: W = 0.94, P = 0.57; island: W = 0.9, P = 0.28) and Mar 07/2018 (mainland: W = 0.96, P = 0.85; island: W = 0.88, P = 0.21) passed the normality test, while timepoint Feb 06/2018 (mainland: W = 0.52, P < 0.001; island: W = 0.73, P = 0.004) failed the normality test. All statistical analyses were performed using R version 4.0.2 and 4.0.5 (R Core Team 2020, 2021).

### Simulating variation captured in the ex situ collection using seed morphological traits

For each of the 14 measured and derived seed traits, we conducted a separate simulation quantifying morphological variation captured when increasing the number of maternal families sampled from contemporary Torrey pine seed collections. Simulations were conducted in R version 3.6.3 (R Core Team 2020) using a customized script [See Supporting Information Figure S2]. Resampling of ex situ collections were performed for island and mainland Torrey pine populations independently, using between one and the total number of maternal families available within each ex situ population collection (mainland: 80 maternal families, island: 30 maternal families) (N_fam_). Maternal trees were sampled randomly without replacement from the pool of available families. All seeds within each selected maternal family were sampled as part of this simulation, except those with missing values for the trait simulated. Overall, between two to five seeds per maternal family were sampled within each population.

To evaluate the number of maternal families needed to capture 95% of seed trait variation in both island and mainland populations, we estimated the number of unique seed trait values captured in a sample of N_fam_ maternal families (N_c_) relative to the total number of unique seed trait values present in a seed population (N_t_). Here, we define “unique seed trait values” as the number of non-redundant standardized measurements for the seed trait simulated rounded to the first digit. Seed morphological measurements were rounded to the first digits as we believe that seed trait variation estimated using additional digits is more likely to fail to capture meaningful biological variation. Standardization of the data was performed so that all seed traits share the same unit (the number of standard deviations a value is from the overall trait mean across populations, see equation (1) above) and become comparable. Sampling of maternal families and estimation of the summary statistic, defined as the proportion of total seed trait variance captured (N_c_/N_t_), were repeated 500 times for each seed morphological trait and Torrey pine population. In this way, N_c_/N_t_ accounts for potential variation in number of seeds sampled per maternal family or variation in sampled maternal families included.

Finally, for each number of maternal trees sampled (N_fam_), we averaged the summary statistic across all 500 replicates. This process was repeated for each of the 14 seed morphological traits and performed for each Torrey pine population separately. Following this, the summary statistic was averaged across all seed traits and separated by populations (see Results below). Proportions of total seed trait variance captured (N_c_/N_t_) are provided based on proportions of maternal families sampled (instead of the number of maternal families sampled) as sample sizes varied across Torrey pine populations.

## Results

### Island-mainland differentiation in seed morphology

A principal component analysis (PCA) using all 14 measured and derived seed traits averaged by maternal family revealed substantial differences in seed morphology between island and mainland populations of Torrey pine (Fig. 3). The first PC axis explained 57.8% of variation in seed morphological traits, primarily separating the island from the mainland population. Seed length, seed width, seed area, endosperm area, and seed coat area exhibited the five highest loadings (absolute values) on PC1 [See Supporting Information Table S1], indicating that seed size and seed coat thickness can largely discriminate island from mainland individuals. On average, seeds collected on island trees were longer, wider, larger, and thicker than seeds collected on mainland trees (Table 1). The second PC axis explained 15.9% of seed trait variation and summarizes within population variability in seed morphology (Fig. 3). Relative seed coat size, relative endosperm size, and relative embryo size had the three highest loadings (absolute values) on PC2 [See Supporting Information Table S1]. This suggests that once corrected for seed size, seed coat thickness, endosperm size, and embryo size are traits contributing to within-population variation.

**Fig. 3.**
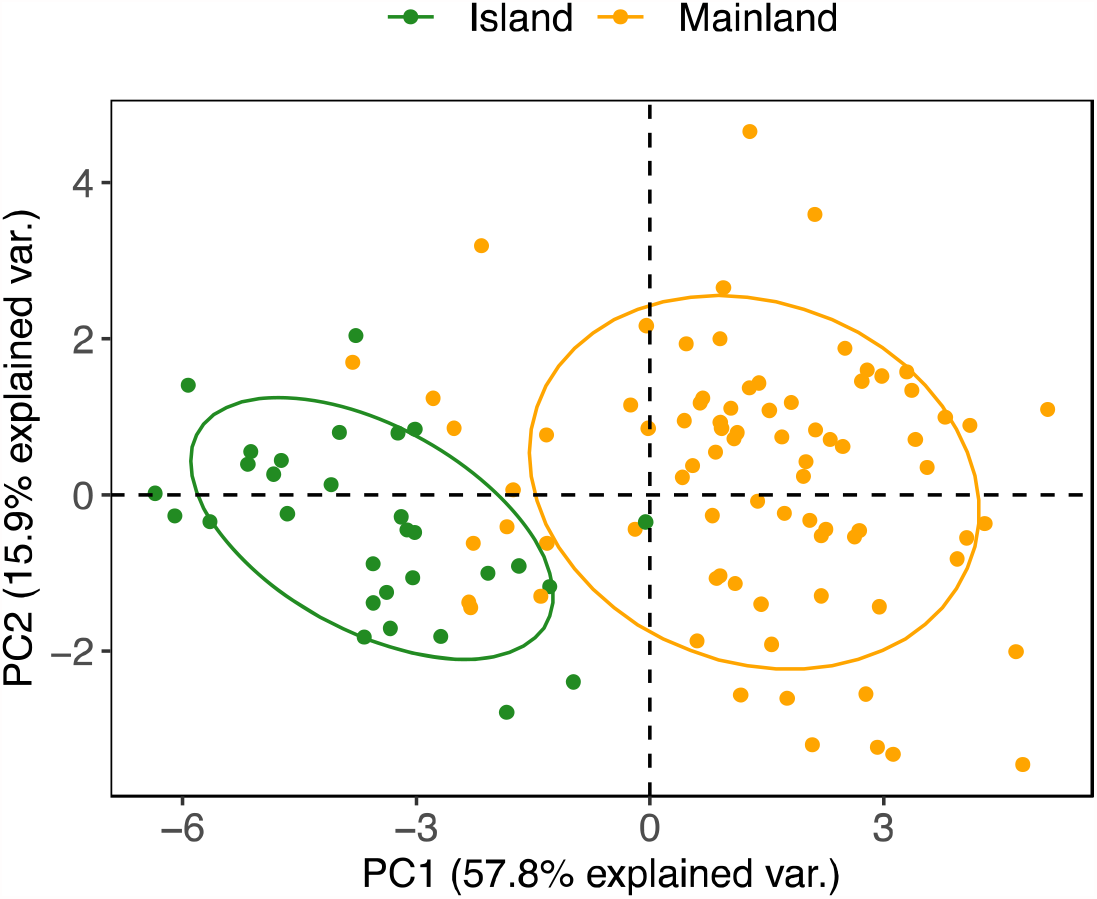
Principal components analysis (PCA) using all 14 seed morphological traits measured and derived from maternal plants collected on Santa Rosa Island (green) and at the Torrey Pine State Reserve (orange).

### Contribution of population origin and maternal family to seed trait variation

Consistent with our principal component analysis, linear mixed models constructed for each of the 14 measured and derived seed traits demonstrated that considerable variation in seed morphology in Torrey pine is explained by population origin (Fig. 4). On average, population origin explained 23% (range = 0.02–0.57) of variation across the species’ distribution [See Supporting Information Table S2]. Traits associated with seed size and seed coat thickness exhibited the highest proportion of variance explained by population origin. These include seed coat area (0.57; F_1,107.60_ = 221.91, P < 0.001), seed area (0.49; F_1,107.56_ = 156.45, P < 0.001), endosperm area (0.37; F_1,108.07_ = 100.58, P < 0.001), seed width (0.36; F_1,108_ = 126.04, P < 0.001), seed coat width (0.32; F_1,108_ = 96.04, P < 0.001), and seed length (0.30; F_1,108.50_ = 78.92, P < 0.001). Overall, this suggests seed size and seed coat thickness are major discriminants of island and mainland Torrey pine seeds.

**Fig. 4.**
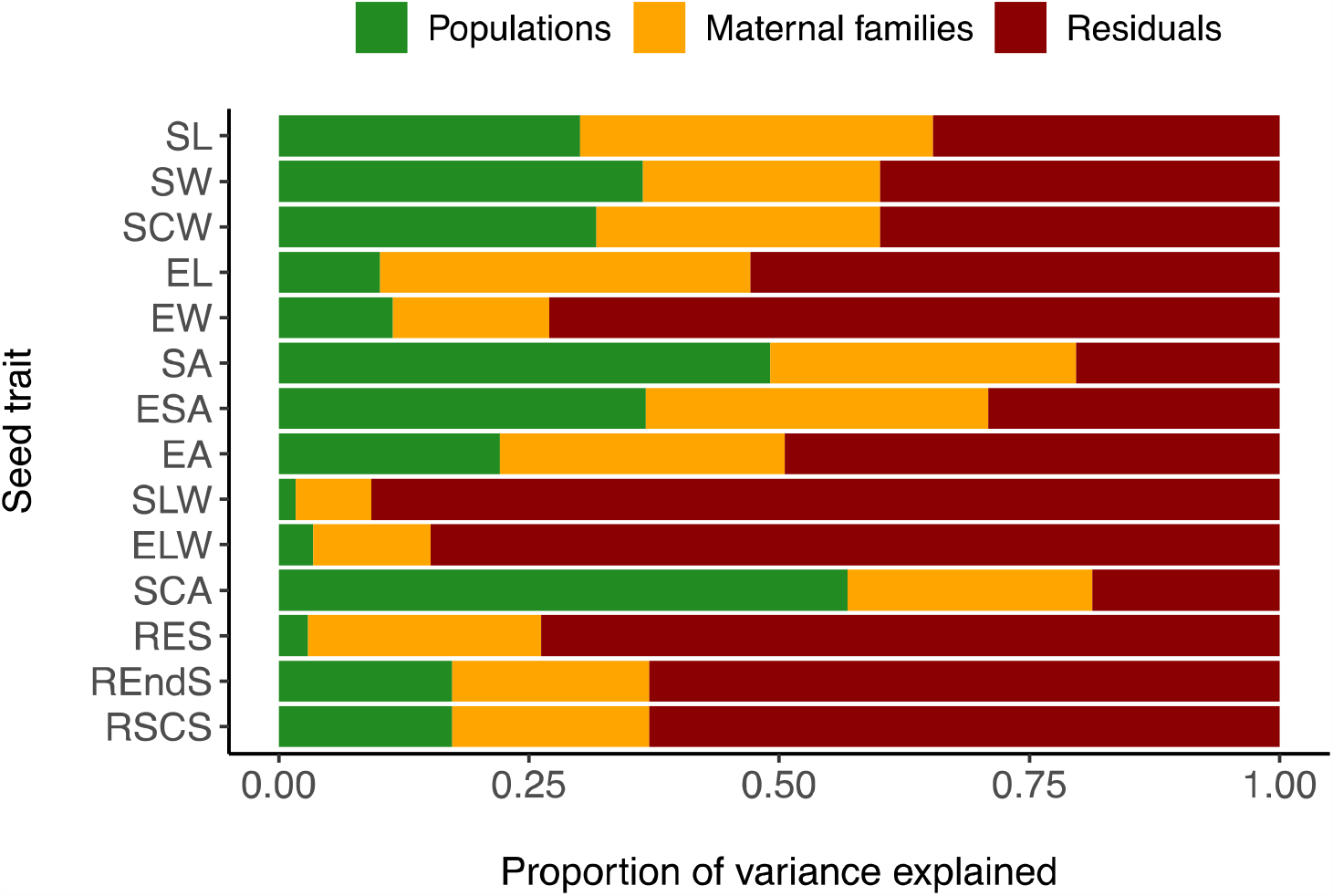
Proportion of variance in seed morphology explained by populations (green), maternal families within populations (orange), and other variables not accounted for in the model (residuals; dark red) for each of the 14 measured and derived seed traits. SW, seed width (cm); SLW, seed length/width ratio; SL, seed length (cm); SCW, seed coat width (cm); SCA, seed coat area (cm^2^); SA, seed area (cm^2^); RSCS, relative seed coat size; RES, relative embryo size; REndS, relative endosperm size; EW, embryo width (cm); ESA, endosperm area (cm^2^); ELW, embryo length/width ratio; EL, embryo length (cm); EA, embryo area (cm^2^). See Supporting Information Table S2 for numerical estimates.

While population origin explained substantial variation across populations, assessment of maternal seed families within populations indicated substantial family structure to seed trait variation (Fig. 4). On average, maternal seed family explained 24% (range = 0.07–0.37) of variation within populations [See Supporting Information Table S2]. Embryo length (0.37; χ ^2^ = 124.82, df = 1, P < 0.001), seed length (0.35; χ ^2^ = 180.72, df = 1, P < 0.001), endosperm area (0.34; χ ^2^ = 211.87, df = 1, P < 0.001), seed area (0.31; χ ^2^ = 256.71, df = 1, P < 0.001), embryo area (0.29; χ ^2^ = 100.16, df = 1, P < 0.001), and seed coat width (0.28; χ ^2^ = 126.93, df = 1, P < 0.001) exhibited the highest proportion of seed trait variation explained by within-population maternal families. This suggests that there is substantial family-level structure to seed size, endosperm size, embryo size, and seed coat thickness within Torrey pine populations.

### Impact of population seed trait differentiation on seedling emergence

The proportion of emerged seedlings increased over time for both island (Q = 15.5, df = 2, P < 0.001) and mainland (Q = 15.2, df = 2, P < 0.001) populations (Fig. 5). However, we found no significant differences in the proportion of individuals emerging between populations across observed time points. On average, 7% and 9% of mainland and island seedlings emerged a month after sowing (Feb 06/2018; W = 28, P = 0.64), 63% and 53% of mainland and island seedlings emerged a month and a half after sowing (Feb 16/2018; t = 0.81, df = 14, P = 0.43), and 78% of mainland and island seedlings emerged two months after sowing (Mar 07/2018; t = -0.06, df = 14, P = 0.95). Overall, this indicates that under controlled conditions, timing and probability of emergence may not be impacted by population differences in seed morphology for Torrey pine seedlings.

**Fig. 5.**
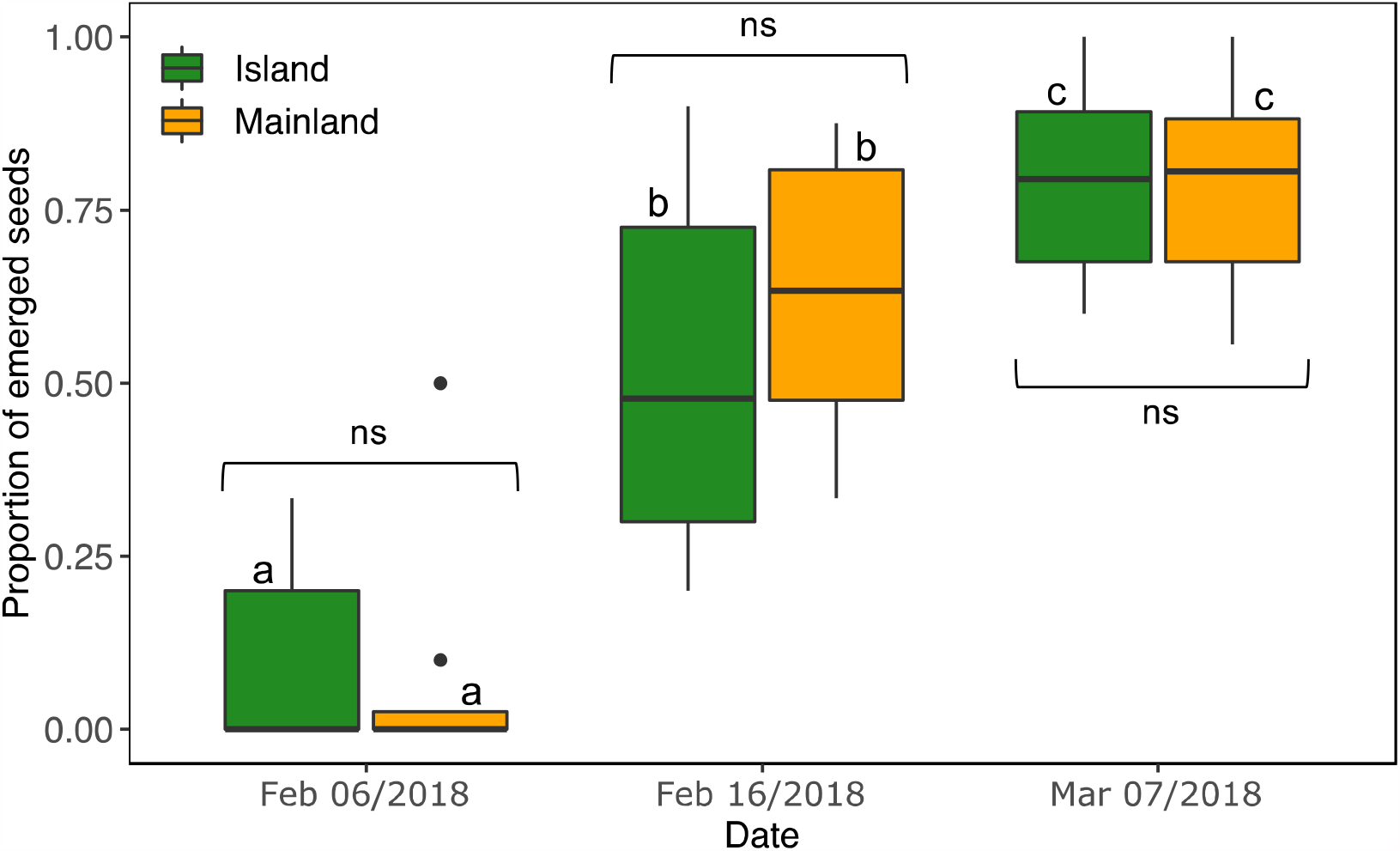
Proportion of emerged seedlings (y axis) recorded at three different timepoints (x axis) for seeds sampled on Santa Rosa Island (green) and at the Torrey Pine State Reserve (orange). Significant differences in emergence time across timepoints within populations are indicated with different letters. Comparisons between populations at each timepoint is indicated with square brackets. ns, non-significant differences (_α_ =0.05).

### Morphological variation captured in simulated seed collections

Simulations revealed that to capture 95% of seed trait variation present in our existing ex situ collections, on average 83% (25 out of 30) and 71% (57 out of 80) of all island and mainland families would need to be resampled, respectively (Fig. 6). This indicates that both island and mainland populations harbor considerable within-population structure for seed morphological traits. Interestingly, capturing equal morphological variation across seed collections always required a higher proportion of island maternal families to be collected relative to the mainland population.

**Fig. 6.**
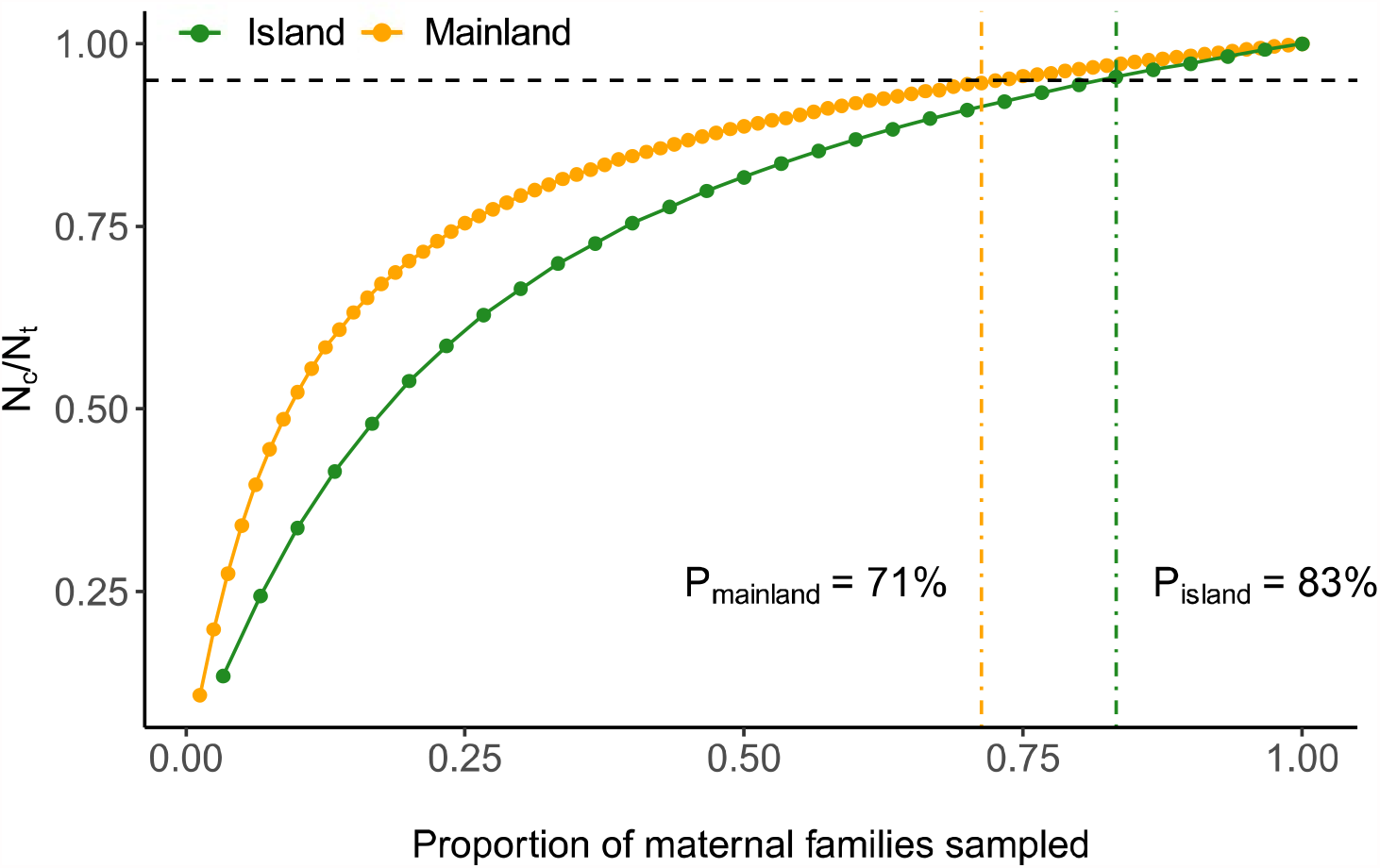
Phenotypic variation captured across seed traits in simulated collections (N_c_) relative to total phenotypic variation present in seed populations (N_t_). Average proportion of phenotypic variation captured (N_c_/N_t_) was estimated for various proportions of maternal families sampled. P_island_ and P_mainland_ represent the proportion of maternal families required to capture 95% of morphological variation (horizontal dashed line) present in island (green) and mainland (orange) ex situ seed populations, respectively.

## Discussion

Evaluating between-and within-population morphological variation in contemporary ex situ seed collections may aid in understanding the distribution of variation needed to guide future conservation efforts. Here, we quantified the distribution of trait variation within an ex situ collection of Torrey pine seeds, with an aim to optimize future supplemental collections. Morphologically, island and mainland seeds were significantly different from each other. Island seeds were larger on average with thicker seed coats relative to their mainland counterparts. These morphological differences may be explained by a combination of stochastic and deterministic factors associated with population origin, including genetic bottlenecks following island colonization, genetic drift, and selection associated with unique biotic pressures. Interestingly, despite substantial morphological differentiation, seedling emergence did not vary among populations, suggesting that either the probability and timing of emergence under controlled conditions is not impacted by differences in seed morphology or that island and mainland seeds respond similarly to an artificial germination protocol. In addition to population origin, a considerable proportion of seed trait variation within each population was explained by maternal family. This suggests that there is substantial within-population variation that will be important to conserve and maintain populations’ evolutionary potential. Finally, simulations demonstrated that 83% and 71% of all maternal families within island and mainland seed collection respectively were necessary to capture 95% of morphological variation within the existing collections. To ensure we maintain representative levels of trait variation in future seed collections, our results suggest that the number of maternal families sampled within natural populations should be maximized, with the island population potentially requiring additional sampling efforts.

Species distributed between island and mainland origins often exhibit marked among-population phenotypic differentiation, including differentiation in seed morphology (Burns *et al*. 2012; Lens *et al*. 2013; Kavanagh and Burns 2014; Burns 2016). Our results revealed considerable seed morphological differences between Torrey pine populations (Fig. 3), primarily in seed size and seed coat thickness (Fig. 4; See Supporting Information Table S1). On average, islands seeds were larger and had thicker seed coats than seeds collected on the mainland (Table 1). These results are consistent with previous studies of island-mainland systems that noted island populations exhibited larger seeds relative to mainland populations (Kavanagh and Burns 2014; Burns 2016; Biddick *et al*. 2019). A combination of different factors could contribute to morphological variation among seed populations, including both stochastic and deterministic forces.

On islands, seeds traits associated with long-distance dispersal may be selected against as they can increase the probability an individual would disperse beyond an island’s limits (Cody and Overton 1996; Kavanagh and Burns 2014; Ottaviani *et al*. 2020, but see Burns 2018). For Torrey pine, increased seed size on the island may have evolved to limit potential seed losses via wind-dispersal, as seed mass negatively correlates with dispersal distance in pines (Greene and Johnson 1993; Debain *et al*. 2003, but see Wyse and Hulme 2020). Nonetheless, Torrey pine seeds possess degenerated wings (Ledig and Conkle 1983), suggesting that other mechanisms likely contribute to seed dispersal in this species. Rodents and birds both feed on Torrey pine, suggesting that seeds may undergo animal-mediated dispersal (Johnson *et al*. 2003). Thus, seed predation may contribute to differences in seed size observed between populations. On the island, *Peromyscus maniculatus* (Deer mouse) is the only rodent present to predate on Torrey pine seeds (Johnson *et al*. 2003). This contrasts with the mainland, where multiple seed predators have been documented; including *Peromyscus boylei* (Brush mice), *Peromyscus maniculatus* (Deer mice), *Peromyscus eremicus* (Cactus mice), *Chaetodipus californicus* (California pocket mice), *Spermophilus beecheyi* (California ground squirrels), or *Aphelocoma californica* (Scrub jays) (Johnson *et al*. 2003). If large seeds are preferentially targeted by seed predators (Reader 1993; Gómez 2004), reduced seed size on the mainland may have evolved as a consequence of the trade-off between attracting predators to promote seed dispersal and mitigating fitness loss due to seed consumption.

While selection may contribute to population differences, differentiation in seed morphology may result from stochastic evolutionary forces. Founder effects associated with the colonization of Santa Rosa island by mainland individuals, and genetic drift in the face of limited gene flow, may have led to morphological differentiation between Torrey pine populations (Ledig and Conkle 1983). Alternatively, a more complex demographic history of the two populations, including colonization, extinction, and recolonization events may have led to the differences observed between populations (Haller 1986). While both stochastic and deterministic factors may contribute to population differences in seed morphology, additional experiments are required to test mechanistic hypotheses. Seeds evaluated in this manuscript were collected from natural populations. To tease apart the contribution of environment and genetics to seed trait differences observed among populations, a common garden experiment is required. Indeed, a reciprocal transplant experiment would be the most effective test of the action of natural selection in shaping morphological differences between island and mainland seeds.

Despite significant differences in seed morphology between populations, timing and probability of emergence was similar across populations (Fig. 5). Emergence rates were high throughout the trial, with 78% of island and mainland seedlings emerging two months after sowing. The absence of differences in seedling emergence between populations was surprising, as seed size often negatively correlates with time to germination (Daws *et al*. 2005; Tanveer *et al*. 2013). However, seed coat thickness can also influence rates of emergence. Seeds with thick seed coats relative to their mass often germinate later than seeds with thinner seed coats (Daws *et al*. 2005; Hamilton *et al*. 2013). For Torrey pine, Hamilton *et al*. (2017) found that island seeds germinate on average two days after mainland seeds. Interestingly, island seeds were not only larger, but also had thicker seed coats relative to mainland seeds (Table 1). Even after correcting for differences in seed size, seed coat thickness (relative seed coat size; RSCS) remained moderately higher in island seeds. Together, these results predict that island seedlings should emerge at similar or later timepoints relative to mainland seedlings, which is consistent with current and previous observations.

Similar emergence rates may also result from our experimental design. Abe and Matsunaga (2011), in a mainland-island comparison study, observed that cold stratification attenuates differences in germination rates between populations of *Rhaphiolepis umbellata*. Additionally, complete and rapid germination of pine seeds is generally observed when pretreated under cold and moist conditions (Krugman and Jenkinson 2008). Overall, this suggests that cold stratification may mask population-specific differences in seedling emergence. Concurrent seedling emergence from both Torrey pine populations coupled with high emergence success suggests a cold stratification protocol is valuable for Torrey pine, particularly where simultaneous emergence for nursery-grown seedlings is desired. Note, however, that variation in the proportion of emerged seedlings within populations across timepoints may have concealed population-specific differences in emergence rates. Consequently, weak differences in the timing and probability of seedling emergence observed between island and mainland populations may be an artifact of small numbers of seeds and maternal families used during emergence trials.

Although population origin explained a substantial proportion of seed trait variation, linear mixed models demonstrated that maternal seed families within populations explained as much variation (Fig. 4; See Supporting Information Table S2). Given generally high heritability for seed morphological traits and the half-sib design of our collection (Pandey *et al*. 1994; Cober *et al*. 1997; Mera *et al*. 2004; Roy *et al*. 2004; Carles *et al*. 2009; Zas and Sampedro 2015; Hakim and Suyamto 2017), family-level seed trait variation likely provides a useful proxy for assessing within-population genetic diversity. With nearly 25% of variation explained on average by maternal families [See Supporting Information Table S2], this suggest there is substantial genetic structure within Torrey pine populations. These results were notable as previous studies using allozymes and chloroplast DNA suggested that the species exhibits little to no within-population genetic variability (Ledig and Conkle 1983; Waters and Schaal 1991; Whittall *et al*. 2010). However, the common garden experiment conducted by Hamilton and colleagues (2017) indicated substantial family-level variation in tree height within both island and mainland populations. Overall, these results indicate that Torrey pine populations may possess within-population genetic variation necessary for natural selection to act upon. From a conservation perspective, these findings suggest that a strategy maximizing the number of maternal families sampled would optimize genetic diversity preserved in future ex situ seed collections and increased distance among individuals may limit relatedness among maternal trees.

Generally, ex situ seed collections aim to capture 95% of genetic diversity present throughout a species’ distribution (Marshall and Brown 1975; Brown and Marshall 1995; Li *et al*. 2002; Gapare *et al*. 2008). Simulations revealed that, in order to capture 95% of morphological variation currently maintained ex situ, 25 (83% of island collection) and 57 (71% of mainland collection) maternal families within each seed collection would need to be sampled (Fig. 6). These data indicate that sampling more maternal families from the island population may be necessary to achieve the same level of representation of morphological variation. Assuming increased phenotypic variation observed on the island results from higher allelic diversity, capturing 95% of genetic variation within the island population will always require more maternal families relative to the mainland population. For these simulations, we assumed that contemporary ex situ collections captured all morphological variation both within and between populations, including seed phenotype frequencies. However, if this is not the case, these recommendations may result in suboptimal sampling of standing variation within targeted populations. This caveat is important because the number of x-rayed maternal families differed between island (30 maternal families) and mainland (80 maternal families) seed collections. To address this caveat, it will be important to have a general understanding of the fraction of natural morphological variation captured across ex situ seed populations and adapt sampling efforts accordingly.

Practical and cost-effective, long-term storage of seeds ex situ is widely used to capture and maintain rare species genetic diversity. These seed collections represent an invaluable resource to quantify within and between population trait variation that may be used to guide future ex situ sampling efforts. Using Torrey pine as a model, we demonstrate that incorporating existing information from ex situ collections offers a unique opportunity to monitor and optimize conservation objectives, particularly important for rare species. While our results and recommendations may be specific to Torrey pine, the empirical, statistical, and simulation-based approaches presented here are broadly applicable to heritable traits across ex situ seed collections.

## Data

The data for this article, including seed morphological measurements and R scripts used are available from GitHub: https://github.com/lnds-anonymous/AoBP2021.

## Acknowledgements

The authors thank Annette Delfino Mix, Valerie Gallup, Andrew Bower, Drew Peterson, Emma Ordemann, Conner Harrington, Forest Swaciak, Jill Wulf, and Stephen Johnson for their help collecting the Torrey pine cones. The collections were funded by The USDA Forest Service, State and Private Forestry, Gene Conservation program to J.W.W. and J.A.H. Sara Wilson provided training and assistance for the use of the x-ray machine. We also thank the USDA Forest Service (Pacific Southwest Research Station) for their support of this project, as well as the Michumash and the Kumeyaay people as the traditional caretakers of the Torrey pine ecosystems sampled for this study. We acknowledge a fellowship to L.N.D.S from the Morton Arboretum Center for Tree Science and thank Sean Hoban for helpful discussion. This work was also supported by a new faculty award from the office of the North Dakota Experimental Program to Stimulate Competitive Research (ND-EPSCoR NSF-IIA-1355466) and funding from the NDSU Environmental and Conservation Sciences Program to J.A.H. Any use of product names is for information purposes only and does not imply endorsement by the US Government.

