## Supporting Information for "Seed morphological traits as a tool to quantify variation maintained in *ex situ* collections: a case study in *Pinus torreyana* (Parry)"

**Table S1.** PC1 and PC2 loadings for all 14 measured and derived seed traits: seed length (SL, cm), seed width (SW, cm), embryo length (EL, cm), embryo width (EW, cm), seed coat width (SCW, cm), seed area (SA, cm^2^), endosperm area (ESA, cm^2^), embryo area (EA, cm^2^), seed length/width ratio (SLW), embryo length/width ratio (ELW), relative embryo size (RES), relative endosperm size (REndS), seed coat area (SCA, cm^2^), and relative seed coat size (RSCS).

| **Seed trait** | **PC1 –57.8% var. explained** | **PC2 –15.9% var. explained** |
| --- | --- | --- |
| SL | -0.33 | -0.06 |
| SW | -0.34 | 0.02 |
| SCW | -0.29 | 0.16 |
| EL | -0.27 | 0.17 |
| EW | -0.27 | 0.31 |
| SA | -0.34 | -0.09 |
| ESA | -0.33 | -0.01 |
| EA | -0.31 | 0.23 |
| SLW | 0.11 | -0.15 |
| ELW | 0.13 | -0.33 |
| SCA | -0.34 | -0.18 |
| RES | 0.09 | 0.46 |
| REndS | 0.18 | 0.45 |
| RSCS | -0.18 | -0.45 |

**Table S2.** Proportion of variance in measured and derived seed morphological traits explained by populations (TPSR, SRI; fixed effect), maternal families (random effect), and both populations and maternal families. SL, seed length (cm); SW, seed width (cm); EL, embryo length (cm); EW, embryo width (cm); SCW, seed coat width (cm); SA, seed area (cm^2^); ESA, endosperm area (cm^2^); EA, embryo area (cm^2^); SLW, seed length/width ratio; ELW, embryo length/width ratio; RES, relative embryo size; REndS, relative endosperm size; SCA, seed coat area (cm^2^); RSCS, relative seed coat size; *R*^2^_m_, marginal variance explained by fixed effect; *R*^2^_c_, conditional variance explained by fixed and random effects.

| **Seed trait** | | **Variance explained by populations**  **(*R*^2^_m_) ^a^** | | **Variance explained by maternal families within populations**  **(*R*^2^_c_-*R*^2^_m_) ^a^** | | **Total variance explained**  **(*R*^2^_c_)** |
| --- | --- | --- | --- | --- | --- | --- |
| SL | 0.30 | | 0.35 | | 0.65 | |
| SW | 0.36 | | 0.24 | | 0.60 | |
| SCW | 0.32 | | 0.28 | | 0.60 | |
| EL | 0.10 | | 0.37 | | 0.47 | |
| EW | 0.11 | | 0.16 | | 0.27 | |
| SA | 0.49 | | 0.31 | | 0.80 | |
| ESA | 0.37 | | 0.34 | | 0.71 | |
| EA | 0.22 | | 0.29 | | 0.51 | |
| SLW | 0.02 | | 0.07 | | 0.09 | |
| ELW | 0.03 | | 0.12 | | 0.15 | |
| SCA | 0.57 | | 0.24 | | 0.81 | |
| RES | 0.03 | | 0.23 | | 0.26 | |
| REndS | 0.17 | | 0.20 | | 0.37 | |
| RSCS | 0.17 | | 0.20 | | 0.37 | |
| *Average* | *0.23* | | *0.24* | | *0.48* | |

^a^ The effect of population origin or maternal family within populations is significant at α=0.05 for all seed traits listed.

**Figure S1.** Torrey pine distribution (red shades) and position of maternal trees collected for cones (closed black circles). (A) Torrey Pine State Reserve, *Pinus torreyana subsp torreyana*. (B) Santa Rosa Island, *Pinus torreyana subsp insularis.*

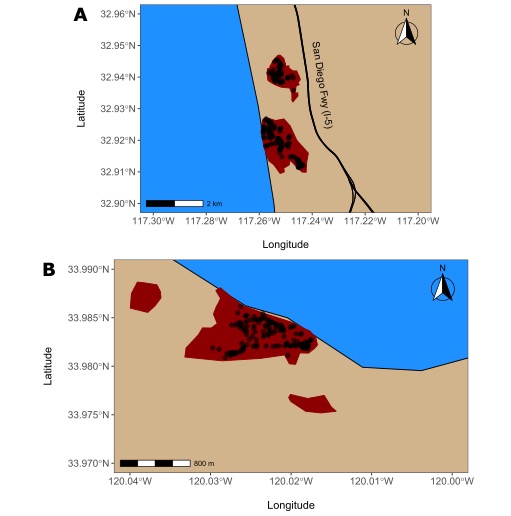

**Figure S2.** Simulation framework used to estimate phenotypic variation in seed morphology captured after resampling of contemporary Torrey pine *ex situ* seed collections. Simulations using this framework were conducted for each seed traits and Torrey pine population independently. Computation proceeds from top to bottom.

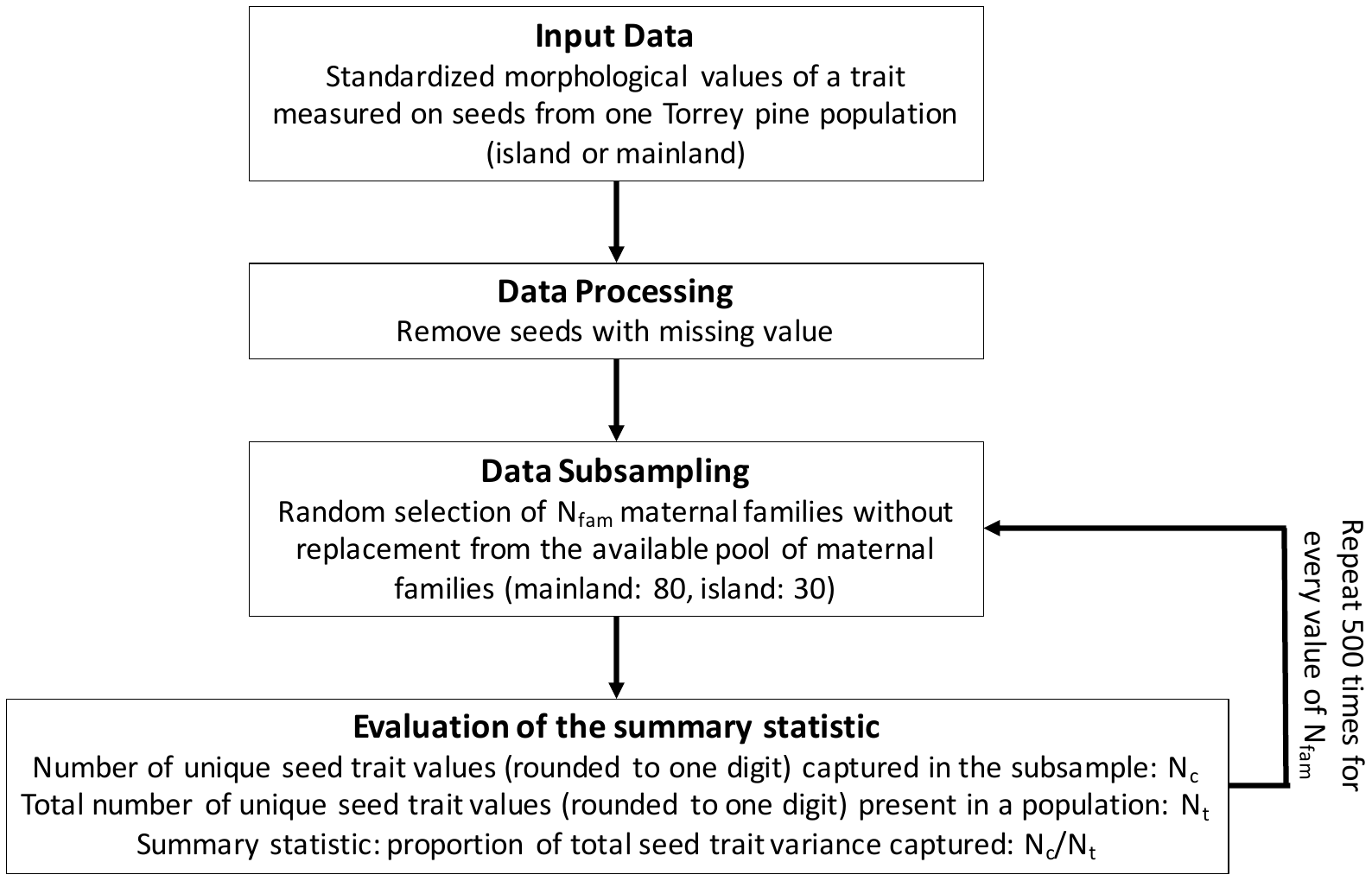
